# Effects of vapor inhalation of 6-methyl nicotine in female and male rats

**DOI:** 10.64898/2026.08.20.746016

**Authors:** Michael A. Taffe, Helen S. Kim, Tess A. Doran, Tyra R. Coons, Sara R.M.U. Rahman, Yanabel Grant, Sophia A. Vandewater

**Author notes:** Address Correspondence to: Dr. Michael A. Taffe, Department of Psychiatry, 9500 Gilman Drive; University of California, San Diego, La Jolla, CA 92093; USA. equal contribution (Tess A. Doran, Tyra R. Coons).

## Abstract

**Background:** The nicotine analog 6-methyl nicotine (6-MN) has appeared in commercial e-cigarette liquids, and other products, spurring interest in determining the extent to which it conveys similar effects to those of nicotine.

**Objective:** To determine if 6-MN acts like nicotine to decrease body temperature, decrease nociception, suppress wheel activity and reinforce operant behavior when delivered by vapor inhalation using an Electronic Nicotine Delivery System (ENDS; “e-cigarette”) approach in a rat model.

**Methods:** Male and female (N=8 per sex) young adult Sprague-Dawley rats were evaluated for rectal temperature and nociceptive responses (warm water tail-withdrawal) to the inhalation of vapor from (-)-6-MN or (-)-nicotine in concentrations ranging from 5-30 mg/mL in the propylene glycol vehicle. Rats were then assessed for the reinforcing effects of nicotine and 6-MN using a vapor self-administration procedure and the rate suppressing effects of nicotine and 6-MN on wheel activity following injection.

**Results:** Inhalation of nicotine or 6-MN for 30 minutes decreased the rectal temperature and increased tail-withdrawal latency of female and male rats in a concentration-dependent manner. The magnitude of the effects of 6-MN and nicotine were similar at similar vapor concentrations. Operant responding for 6-MN vapor was increased by pre-treatment with the antagonist mecamylamine. 6-MN was more potent than nicotine at suppressing wheel activity after injection.

**Conclusions:** 6-MN induces effects very similar to those of nicotine, at a similar potency when inhaled and at a slightly increased potency when injected.

## 1. Introduction

The 6-methyl nicotine (6-MN) analog of nicotine occurs in tobacco products, but in an abundance ~4 orders of magnitude less than nicotine content in tobacco and traditional nicotine e-liquids (Pankow et al., 2025). In recent years, however, the 6-MN analog has been confirmed as the primary psychoactive drug in some liquids marketed for use in Electronic Nicotine Delivery Systems (ENDS), as well as in oral pouch products (Jordt and Jabba, 2024; Jordt et al., 2024; Mallock et al., 2024; Vanhee et al., 2024). The 6-MN analog also appears to be the active ingredient in a new combustible, non-tobacco cigarette product (Seidenberg et al., 2026). Studies reported a concentration of 6-MN in several e-liquid products available in the USA that was ~10% or less of the labeled concentration of 50 mg/g (Erythropel et al., 2024; O’Connor et al., 2026), and another found concentrations 20-30% of the labeled concentration in products in Australia (Jenkins et al., 2024). This potentially suggests a market-based interpretation of increased potency of 6-MN compared to nicotine, and therefore marketing intended to signal this to the consumer.

Well controlled laboratory reports which examine the relative potency of 6-MN and nicotine are scarce, as reviewed (Effah et al., 2025), although it has been reported that (+/-)-6MN is about 3-fold more potent than (+/-)-nicotine in tail-withdrawal anti-nociception and 5-fold more potent in inhibiting spontaneous activity in mice (Dukat et al., 2002). It has also been reported that 6-MN is behaviorally equipotent with nicotine using a drug-discrimination assay in male rats (Levy and Dunn, 1979). Although most prior *in vivo* investigation of nicotine effects used parenteral injection, recent developments show it is also possible to determine the effects of nicotine delivered by ENDS vapor inhalation (Espinoza et al., 2022; Javadi-Paydar et al., 2019; Javadi-Paydar et al., 2024; Montanari et al., 2020), including as a reinforcer of operant responding in rats (Gutierrez et al., 2024a; Gutierrez et al., 2024b; Lallai et al., 2021; Smith et al., 2020) and mice (Cooper et al., 2021; Henderson and Cooper, 2021). For example, acute non-contingent administration of (-)-nicotine ditartrate by vapor inhalation reduces core body temperature of rats and can increase spontaneous locomotor behavior under some conditions (Javadi-Paydar et al., 2018; Javadi-Paydar et al., 2019) and discontinuation from repeated exposure induces somatic signs of withdrawal (Martinez et al., 2023; Montanari et al., 2020).

We recently reported nicotine and 6-MN to be approximately equipotent in a limited investigation of temperature, nociception and suppression of wheel activity in nicotine-experienced middle aged female Wistar rats (Taffe et al., 2026). The present study was designed to extend that study by examining the impact of 6-MN inhalation in young adult animals without extensive prior nicotine exposure and to determine sex effects by comparing the impact of 6-MN in male and female rats.

## 2. Methods

### 2.1 Subjects

Male and female (N=16 per sex) Sprague-Dawley (Envigo) rats, received as two Cohorts in the laboratory at 11-15 weeks of age, were used for this study. The experimental drug exposures were initiated at 18 and 17 weeks of age for Cohort 1 and 2 respectively. The vivarium was kept on a 12:12 hour reversed light-dark cycle, and behavior studies were conducted during the vivarium dark period. Food and water were provided *ad libitum* in the home cage. Procedures were conducted in accordance with protocols approved by the Institutional Animal Care and Use Committee of the University of California, San Diego and were consistent with the NIH Guide (Garber et al., 2011).

### 2.2 Drugs

(-)-Nicotine ditartrate (Sigma-Aldrich, St. Louis, MO) and (-)-6-methyl nicotine (Nicotine River, Thousand Oaks, CA, USA) were dissolved in propylene glycol (PG; Fisher Scientific, Nicotine River) at concentrations of 5, 10, and 30 mg/mL for vapor studies. PG was used as the inhalation vehicle for consistency and comparability with our prior reports on the impact of acute and repeated nicotine vapor inhalation (Gutierrez et al., 2024a; Gutierrez et al., 2024b; Javadi-Paydar et al., 2019). A vehicle of cremulphor:ethanol:saline (1:1:18) was used for i.p. injection of (-)-Nicotine ditartrate (Sigma-Aldrich, St. Louis, MO) and (-)-6-methyl nicotine freebase (LGC, Ltd; Manchester, NH) for the wheel activity experiment.

### 2.3 Apparatus

#### 2.3.1 Vapor Inhalation

Rats were exposed in pairs to 30-minute vapor inhalation sessions for non-contingent exposure. An ENDS based vapor inhalation system (La Jolla Alcohol Research, Inc) was used for these studies, using procedures validated for nicotine exposure (Gutierrez et al., 2024a; Javadi-Paydar et al., 2019; Javadi-Paydar et al., 2024). In brief, vapor was delivered into sealed vapor exposure chambers through the use of controllers which trigger commercial e-cigarette tanks/atomizers (SMOK TFV8 X-baby, 0.25 ohm V8 X-Baby M2 Core; and Geek Vape ZEUS Z Sub-Ohm tanks with 0.25 ohm Z Dual Coils). Vacuum control through an exhaust valve flowed room air at ~1 L per minute and ensured that vapor entered the chamber on each device triggering event. Puffs (6 seconds) were delivered for non-contingent exposure every 5 minutes during the session with the airflow turned on 15 seconds prior to puff initiation and turned off 10 seconds after puff initiation.

### 2.4 Experiments

#### 2.4.1 Experiment 1: Effect of 6-methyl nicotine inhalation on temperature and nociception

For this study, tail withdrawal latency (52°C warm water immersion) and body temperature (rectal thermistor) were obtained prior to the session, using procedures previously described (Gilpin et al., 2011; Nguyen et al., 2018) in groups (N=8 per sex) of male and female rats (Cohort 1). These measures were then re-assessed at 35, and 60 minutes after the start of inhalation. One habituation session of assessment at the three timepoints, without any vapor inhalation, was run four weeks prior (starting on PND91). The impact of (-)-6-methyl nicotine (0, 5, 10 and 30 mg/mL in the propylene glycol vehicle) was assessed in an order counterbalanced across each pair of animals. Vapor studies were conducted PND 123 – PND 140 with a minimum of three days between evaluations in any given rat.

#### 2.4.2 Experiment 2: Effect of nicotine compared with the effect of 6-methyl nicotine on temperature and nociception

The Cohort 1 rats from Experiment 1 were next evaluated for responses to 30 minutes of inhalation of vapor from PG, (-)-6-methyl nicotine (30 mg/mL) or (-)-nicotine (30 mg/mL) in an order counterbalanced for each pair of animals, starting on PND 147. Tail withdrawal latencies and body temperatures were assessed as in Experiment 1.

Following a vapor self-administration experiment involving 26 sessions of 30 min duration where (-)-6-methyl nicotine (3 mg/mL) or (-)-nicotine (3 mg/mL) puffs were made contingent upon a lever press (see below), these rats were again evaluated for responses to 30 minutes of inhalation of vapor from PG, (-)-6-methyl nicotine (30 mg/mL) or (-)-nicotine (30 mg/mL) in a counterbalanced order starting on PND 214. For these experiments the room temperature was held at 22.5-23.7°C. Room temperature was not consistently monitored in the first evaluation, but in some other experiments was found to be as low as 21.4°C on some days. Ambient temperature can alter responses to drugs (Aarde et al., 2017; Cornish et al., 2003; Malberg and Seiden, 1998; Miller et al., 2013; Von Huben et al., 2007), thus a tighter window was maintained for this follow-up study. The water bath for tail-withdrawal was lowered to 50°C to potentially facilitate detection of relatively weak anti-nociceptive effects observed in the first experiment.

This experiment was repeated in a new group (Cohort 2) of male and female rats (N=8 per sex) to directly contrast the impact of nicotine and 6-MN in drug naïve animals. As in Experiment 1, nociception and body temperature were assessed without any vapor inhalation at the three timepoints in a habituation session on PND 112. Starting on PND 117, the Cohort 2 rats were evaluated for temperature and nociception responses after 30 minutes of inhalation of vapor from PG, (-)-6-MN (30 mg/mL) or (-)-nicotine (30 mg/mL) in an order counter-balanced for each pair of animals. After this the rats were tested after inhalation of (-)-6-MN (60 mg/mL) or (-)-nicotine (60 mg/mL), in an order counterbalanced by pair. Tail withdrawal latencies and body temperatures were assessed as in Experiment 1 with the room temperature maintained at 22.5-23.7°C.

#### 2.4.3 Experiment 3: Mecamylamine antagonism of the effects of nicotine and 6-methyl nicotine inhalation

The Cohort 2 animals were next evaluated following inhalation of PG, 6-MN (60 mg/mL) or nicotine (60 mg/mL) 15 minutes after being injected with either saline or mecamylamine (2 mg/kg, s.c.). The three vapor conditions were evaluated in an order counter-balanced for each pair of animals, with the pre-injection alternated within the pairs. The room temperature was maintained at 22.5-23.7°C for this experiment.

#### 2.4.4 Experiment 4: Vapor self-administration of nicotine and 6-methyl nicotine

The rats were exposed to puffs of 6-MN (3 mg/mL; N=8 per sex) or nicotine (3 mg/mL; N=8 per sex) vapor made contingent upon a lever press (Fixed Ratio 1) in 30-minute sessions with Cohort 1 (N=4 per sex/drug) starting on PND 173 and Cohort 2 (N=4 per sex/drug) starting on PND 197. After 9 sessions the drug identity was switched for another 9 sessions. Thereafter, rats were treated with mecamylamine (0.0, 1.0, 2.0 mg/kg, i.p.) 15 minute prior to the session in a counter-balanced order for first the swapped drug condition and then with their original training drug. Two (Cohort 1) or four (Cohort 2) sessions of their original training drug were run prior to conducting the mecamylamine challenges.

#### 2.4.5 Experiment 5: Effect of nicotine and 6-methyl nicotine on wheel activity

Administration of (-)-nicotine ditartrate or (±)-6MN ditartrate reduced voluntary wheel activity in rats when administered by s.c. injection or vapor inhalation (Gutierrez et al., 2024a; Gutierrez et al., 2024b; Taffe et al., 2026). For this experiment rats were injected 15 minutes prior to being provided wheel access, with activity measured in quarter-rotations every 5 minutes. Studies were conducted starting on PND 269 (Cohort 1) or PND 253 (Cohort 2) with a minimum of three days between evaluations in any given rat. This study in Cohort 1 first evaluated the impact of vehicle (1:1:18 ethanol:cremulphor:saline), (-)-nicotine ditartrate (0.4 mg/kg, s.c.) and (-)-6-MN freebase (0.152 mg/kg, s.c.) in a counter-balanced order. The study was then repeated with vehicle, (-)-nicotine ditartrate (0.2 mg/kg, s.c.) and (-)-6-MN freebase (0.076 mg/kg, s.c.) doses. The Cohort 2 animals completed the lower dose evaluations first and the higher dose evaluations second. Although Cohort 1 animals had received two baseline wheel sessions PND 143-144, there was variable interest in the wheel for the vehicle animals on the first day of this study, thus the Day 1 schedule for Cohort 1 was repeated on a fourth day, which was used for the analysis. Cohort 2 animals received two baseline wheel sessions PND 246-247, i.e., just prior to the drug challenges to minimize any similar novelty effects.

The Cohort 1 animals were assessed for wheel activity following 30-minute inhalation sessions with vapor from PG, (-)-nicotine (30 mg/mL) and (-)-6-methyl nicotine (30 mg/mL), starting on PND 319. For this study, rats were injected with mecamylamine (0, 1, 2 mg/kg, i.p.) immediately prior to the vapor inhalation session.

### 2.6 Data Analysis

Rectal temperature (°Celsius), tail withdrawal latency (seconds), vapor deliveries and wheel activity (quarter rotations) were analyzed by 2-way ANOVA including within-subjects factors for Dose/Concentration, Inhalation, and/or Pretreatment Conditions and Time, and between-subjects factors of sex as relevant. One-way ANOVA was used with a within-subject factor of Dose/Concentration condition to compare the 35-minute timepoint for rectal temperature and tail withdrawal latency data, and the summed wheel activity. Mixed effects analysis was used in any cases where data points were missing.

In all analyses, a criterion of P<0.05 was used to infer a significant difference. Significant main effects were followed with post-hoc analysis using Tukey (multi-level factors), Sidak (two-level factors) or Dunnett (treatments versus control within group) correction. All analysis used Prism for Windows (v. 11.0.x; GraphPad Software, Inc, San Diego CA).

## 3. Results

### 3.1.1 Experiment 1: Effect of 6-methyl nicotine inhalation on temperature and nociception

The inhalation of 6-MN significantly reduced the body temperature, and increased the tail-withdrawal latency of both male and female rats. The decrease in temperature depended on the 6-MN concentration in the female rats and there was a threshold over 5 mg/mL required to increase withdrawal latency in the male rats. The ANOVA confirmed significant effects of Time and Concentration in both male [Concentration: F (2.606, 18.24) = 22.64; P<0.0001; Time: F (1.851, 12.96) = 27.34; P<0.0001] and female [Concentration: F (1.624, 11.37) = 39.53; P<0.0001; Time: F (1.964, 13.75) = 102.6; P<0.0001] rats, and the interaction of factors in female rats [F (2.589, 18.13) = 11.10; P<0.0005] on body temperature. The Tukey post-hoc test limited to orthogonal comparisons confirmed a significant difference between PG and each 6-MN concentration after immediately after inhalation in the female rats and between PG and both 10 and 30 mg/mL 6-MN in both sexes at the 60-minute time-point. Body temperature was significantly lower than the pre-inhalation value for male (5, 30 mg/mL) and female (5, 10, 30 mg/mL) rats immediately after inhalation, as well as at the 60-minute time point for male (5, 30 mg/mL) and female (PG; 10, 30 mg/mL) rats. See **Figure 1** for additional significant differences.

**Figure 1.**
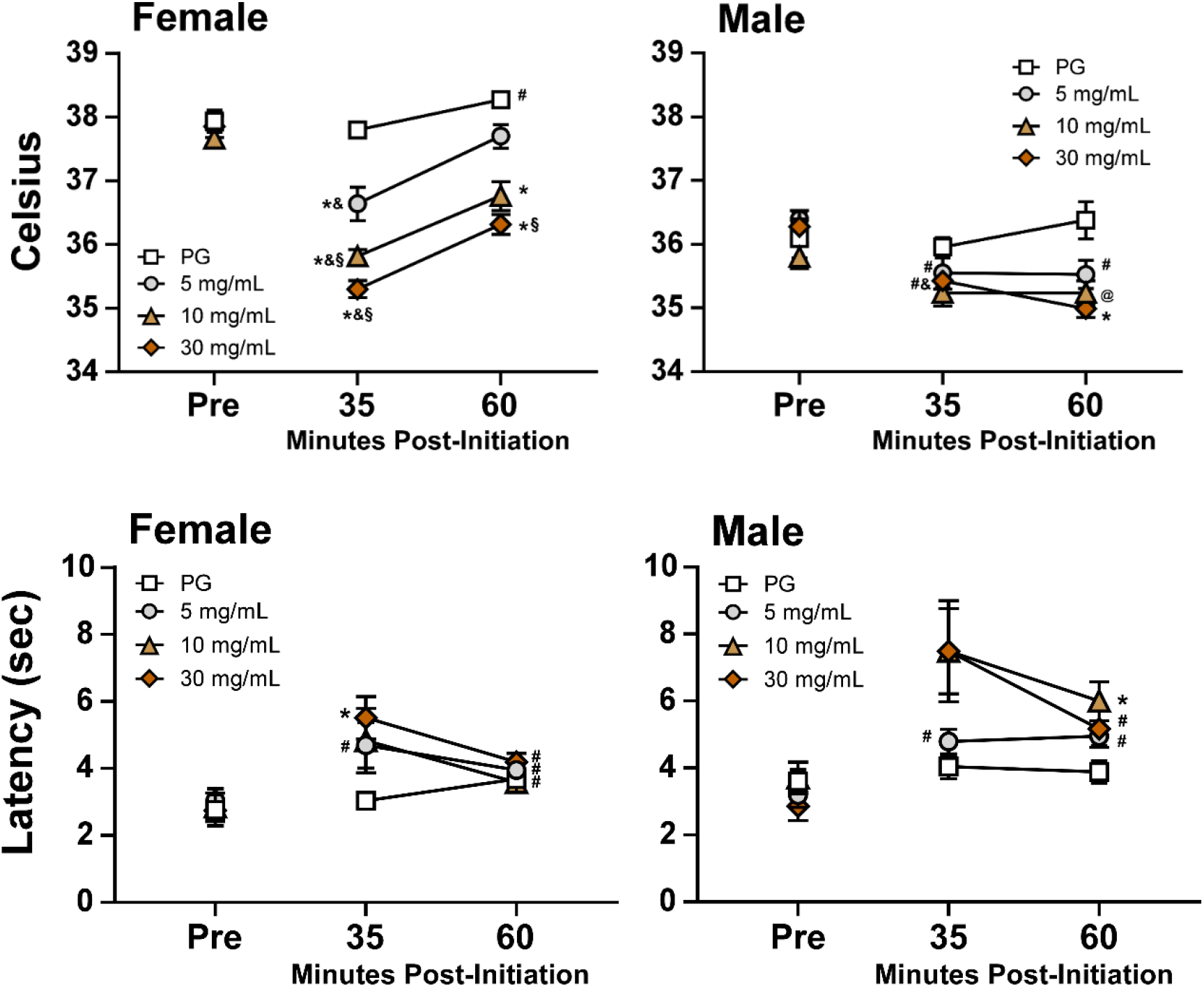
Mean (N=8 per sex; ±SEM) rectal temperature and tail-withdrawal latency assessed before and after inhalation of vapor from the PG vehicle or (-)-6-methyl nicotine (6-MN5, 5, 10, 30 mg/mL in the PG). @ = a significant difference from PG; * = a significant difference from both the pre-treatment value and PG at the respective timepoint; # = a significant difference from the pre-treatment value; & = a significant difference from 60 minute; § = a significant difference from 5 mg/mL; ‡ = a significant sex difference.

The tail-withdrawal latency was modestly affected by 6-MN vapor inhalation. The ANOVA confirmed significant effects on withdrawal latency of Time [F (1.799, 12.59) = 11.03; P<0.005] and Concentration [F (2.016, 14.11) = 7.54; P<0.01] in male rats, and of Time [F (1.313, 9.189) = 20.22; P<0.001] in female rats. The Tukey post-hoc test limited to orthogonal comparisons confirmed a significant difference between PG and 30 mg/mL 6-MN 35 minutes after the start of inhalation in female rats and between PG and 10 mg/mL 6-MN 60 minutes after the start of inhalation in male rats. See **Figure 1** for additional significant differences.

### 3.4.2 Experiment 2: Effect of nicotine and 6-methyl nicotine on temperature and nociception

In the initial direct-comparison study, Cohort 1 female rats’ tail-withdrawal latencies were slowed by vapor inhalation of (-)-6-MN (30 mg/mL) and by (-)-nicotine (30 mg/mL) to a similar extent [Time: F (1.588, 11.12) = 11.55; P<0.005; Interaction of Dose Condition with Time: F (2.561, 17.93) = 4.11; P<0.05]. The Tukey post hoc test confirmed withdrawal latencies were slower 35 minutes after the start of inhalation of either (-)-nicotine or (-)-6-MN compared with PG inhalation (**Figure 2A**). Latencies were also slower relative to baseline at the 35 min timepoint after either (-)-6-MN or (-)-nicotine inhalation and 60 minutes after the initiation of (-)-6-MN inhalation. Withdrawal latency was also faster 60 minutes after the initiation of (-)-6-MN compared to the 35-minute timepoint. The tail-withdrawal was not significantly altered by Time or by Dose condition in male rats, nor was body temperature significantly altered in either sex.

**Figure 2.**
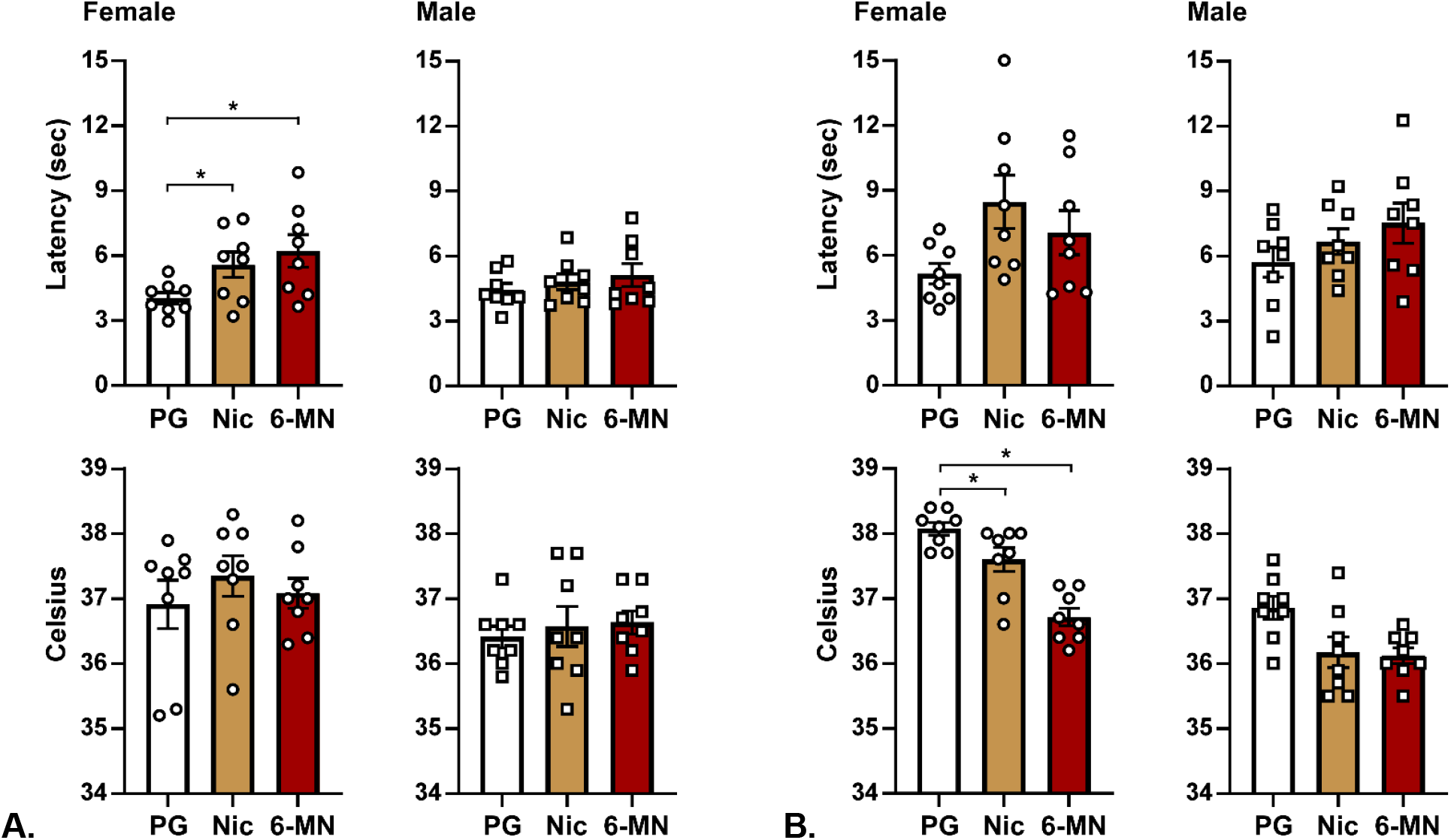
Mean (N=8; ±SEM) and individual tail-withdrawal latency and rectal temperature assessed 35 minutes after the start of inhalation of vapor from the PG vehicle, nicotine (Nic; 30 mg/mL in the PG) or (-)-6-methyl nicotine (6-MN; 30 mg/mL in the PG) starting at **A)** PND 147 and **B)** PND 214 in Cohort 1. Significant differences between inhalation conditions are indicated with *.

The experiment was repeated after conclusion of vapor self-administration experiments. In this case the room temperature was held fixed between 22.5-23.7°C and the water temperature was 50°C. The analysis confirmed a significant effect of vapor condition on a 1 way ANOVA for female temperature, not for male temp (P=0.064) or either tail withdrawal at the 35 minute timepoint (**Figure 2B**). The two-factor ANOVA confirmed a significant effect of vapor condition [F (1.855, 25.97) = 5.69; P<0.05] on tail-withdrawal, but not any effect of sex. Marginal mean post-hoc Tukey confirmed a significant difference between PG and both nicotine (30 mg/mL) and 6-MN (30 mg/mL).

In the two-factor ANOVA, significant effects of vape condition [F (1.544, 21.62) = 15.87; P<0.0005], and of sex [F (1, 14) = 132.7; P<0.0001] on tail-withdrawal were confirmed. In the post-hoc test limited to all orthogonal comparisons, a significant difference between PG and (-)-nicotine (30 mg/mL) and between (-)-nicotine (30 mg/mL) and (-)-6-MN (30 mg/mL) vapor were confirmed for female rats. The post-hoc test also confirmed significant sex differences for each vapor inhalation condition.

#### Cohort 2

The direct contrast experiment was repeated in Cohort 2, drug naïve animals. The contrast of PG, (-)-6-MN (30, 60 mg/mL) and n(-)-icotine (30, 60 mg/mL) in this group confirmed a similar magnitude of effect of (-)-nicotine and (-)-6-MN on rectal temperature and tail-withdrawal latency.

Inhalation of (-)-6-MN decreased body temperature of both female and male rats (**Figure 3A,B**). The ANOVA confirmed that rectal temperatures in the female group were significantly affected by Time [F (1.564, 10.95) = 66.15; P<0.0001] and Inhalation Condition [F (3.023, 21.16) = 5.83; P<0.005] but not by the interaction of Time with Inhalation Condition. The Tukey post-hoc test of all orthogonal comparisons confirmed temperature was lower 35 minutes after initiation of the (-)-6-MN (30, 60 mg/mL) conditions compared with the PG session. Temperature was also significantly lower 35 minutes after the start of inhalation of (-)-Nicotine (30 mg/mL) and (-)-6-MN (30, 60 mg/mL) conditions compared with the pre-treatment temperature, and in all five conditions compared with the 60-minute time point.

**Figure 3:**
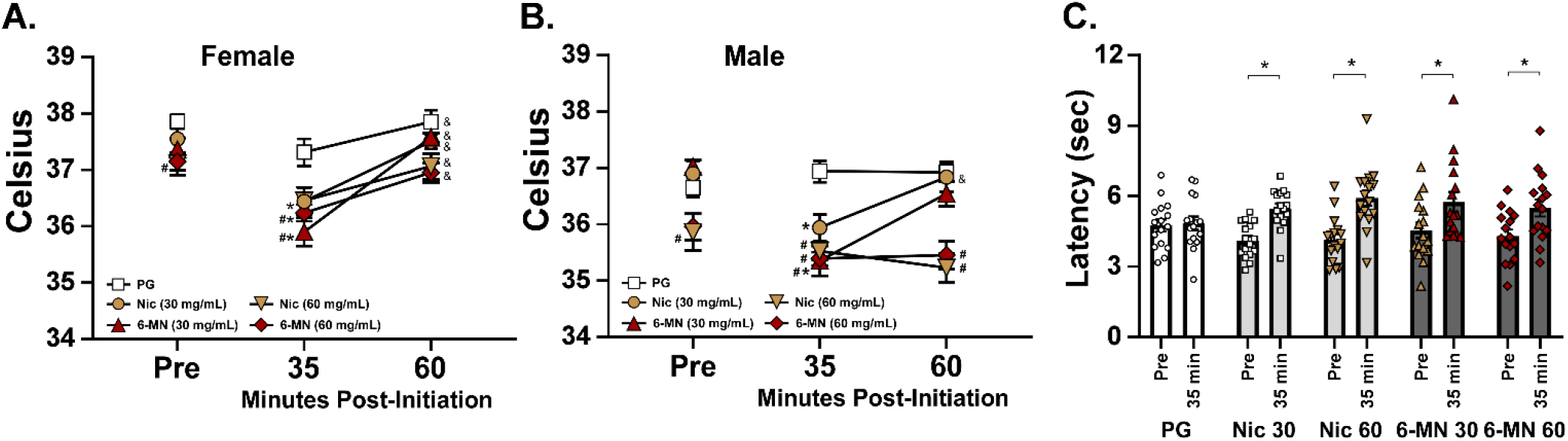
Mean (±SEM) **A)** female (N=8) and **B)** male (N=8) rats rectal temperature, and C) mean and individual tail-withdrawal latencies of both sexes, assessed before and after inhalation of vapor from the PG vehicle, (-)-nicotine (Nic; 30, 60 mg/mL in the PG) or (-)-6-methyl nicotine (6-MN; 5, 30, 60 mg/mL in the PG) for 30 minutes. A significant difference from vehicle vapor (PG) is indicated with #, from the pre-inhalation baseline with * and from the 35 minute timepoint after the start of inhalation with &.

The ANOVA confirmed that rectal temperatures in the male group were significantly affected by Time [F (1.311, 9.178) = 13.83; P<0.005], by Inhalation Condition [F (3.006, 21.04) = 16.21; P<0.0001] and by the interaction of Time with Inhalation Condition [F (3.466, 24.26) = 4.60; P<0.01]. The Tukey post-hoc test of all orthogonal comparisons confirmed temperature was lower 35 minutes after initiation of the Nicotine (60 mg/mL) and 6-MN (30, 60 mg/mL) conditions, and 60 minutes after initiation of the Nicotine (60 mg/mL) and 6-MN (60 mg/mL) conditions, compared with the PG session. Temperature was also significantly lower 35 minutes after the start of inhalation of Nicotine (30 mg/mL) and 6-MN (30 mg/mL), and 60 minutes after the start of Nicotine (60 mg/mL) and 6-MN (60 mg/mL) inhalation, compared with the pre-treatment temperature.

Analysis of all animals, male and female, confirmed a significant effect of Time [F (1.000, 15.00) = 48.74; P<0.0001] and of the interaction of Time with Vape condition [F (3.149, 47.24) = 3.52; P<0.05] on tail withdrawal latency (**Figure 3C**). The Tukey post-hoc test limited to orthogonal comparisons confirmed significantly slower latencies after vaping in all conditions except PG. Follow-up analysis within each sex did not confirm any significant effects of vape condition within male or female, however there was a main effect of Time on tail withdrawal latency in male [F (1.000, 7.000) = 27.16; P<0.005] and female [F (1.000, 7.000) = 20.65; P<0.005] rats and the post-hoc test confirmed a difference from Pre to 35 min in females after nicotine (30 mg/mL) inhalation and in males after nicotine (60 mg/mL) inhalation.

### 3.4.3 Experiment 3: Mecamylamine antagonism of the effects of nicotine and 6-methyl nicotine inhalation

The ANOVA confirmed a significant interaction of Vape Condition with Pre-Treatment [F (1.875, 28.12) = 7.33; P<0.005] on body temperature of all rats (**Figure 4**). The Tukey post-hoc test of all orthogonal comparisons confirmed a difference between the Nicotine inhalation and both other conditions following saline injection and a difference between saline and mecamylamine injection for PG and Nicotine inhalation conditions. This effect of the interaction was also confirmed within the female group [F (1.712, 11.99) = 7.26; P<0.05], but not the male group, however the post-hoc test only confirmed a significant effect of mecamylamine after PG inhalation (not shown).

**Figure 4.**
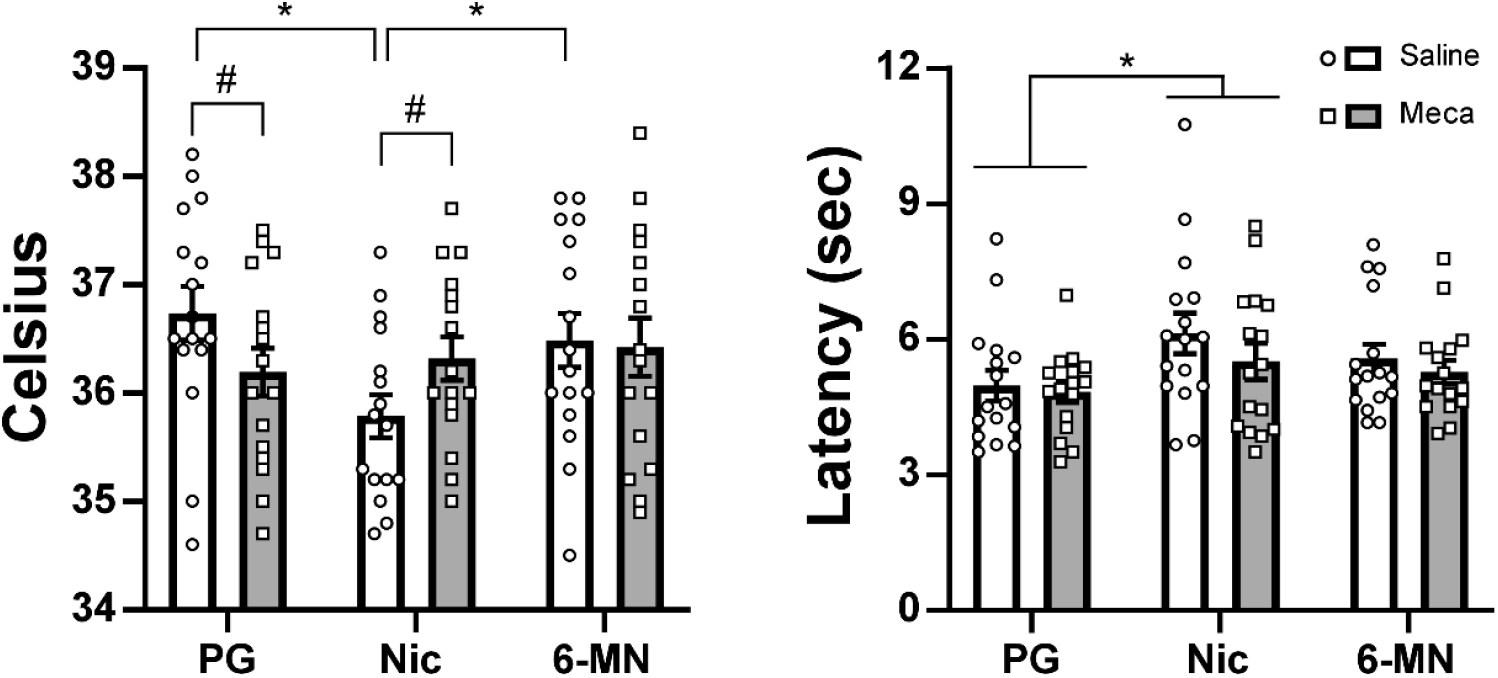
Mean (N=16; ±SEM) and individual rectal temperature and tail-withdrawal latency assessed 35 minutes after the start of inhalation of vapor from the PG vehicle, nicotine (Nic; 60 mg/mL in the PG) or (-)-6-methyl nicotine (6-MN; 60 mg/mL in the PG). A significant difference between Saline and Mecamylamine pre-treatment is indicated with #, and differences between vapor conditions with *.

The ANOVA also confirmed a significant effect of Vape Condition on tail withdrawal latency (**Figure 4**) of all rats [F (1.898, 28.47) = 4.55; P<0.05] and within the male sub-set [F (2, 14) = 4.25; P<0.05]. The Tukey post-hoc test without Greenhouse-Geisser correction confirmed Nicotine increased withdrawal latency compared with PG inhalation in the full sample (**Figure 4**) and the male subset (not shown).

### 3.5 Experiment 4: Vapor self-administration of nicotine and 6-methyl nicotine

The sum of vapor deliveries obtained in the initial acquisition (**Figure 5A**) and the swapped drug sessions (**Figure 5B**) depended on group, with the ANOVA confirming significant effects for the original [Drug: F (1, 28) = 4.91; P<0.05; Interaction: F (1, 28) = 5.06; P<0.05] and swapped [Drug: F (1, 28) = 4.60; P<0.05; Interaction: F (1, 28) = 5.58; P<0.05] sessions. In both cases the Tukey post-hoc test confirmed a significant difference between the female groups. Follow-up analysis confirmed that the males self-administered more vapor deliveries in the switched condition compared with their initial training drug (F (1, 14) = 9.58; P=0.01) but this did not vary by group.

**Figure 5:**
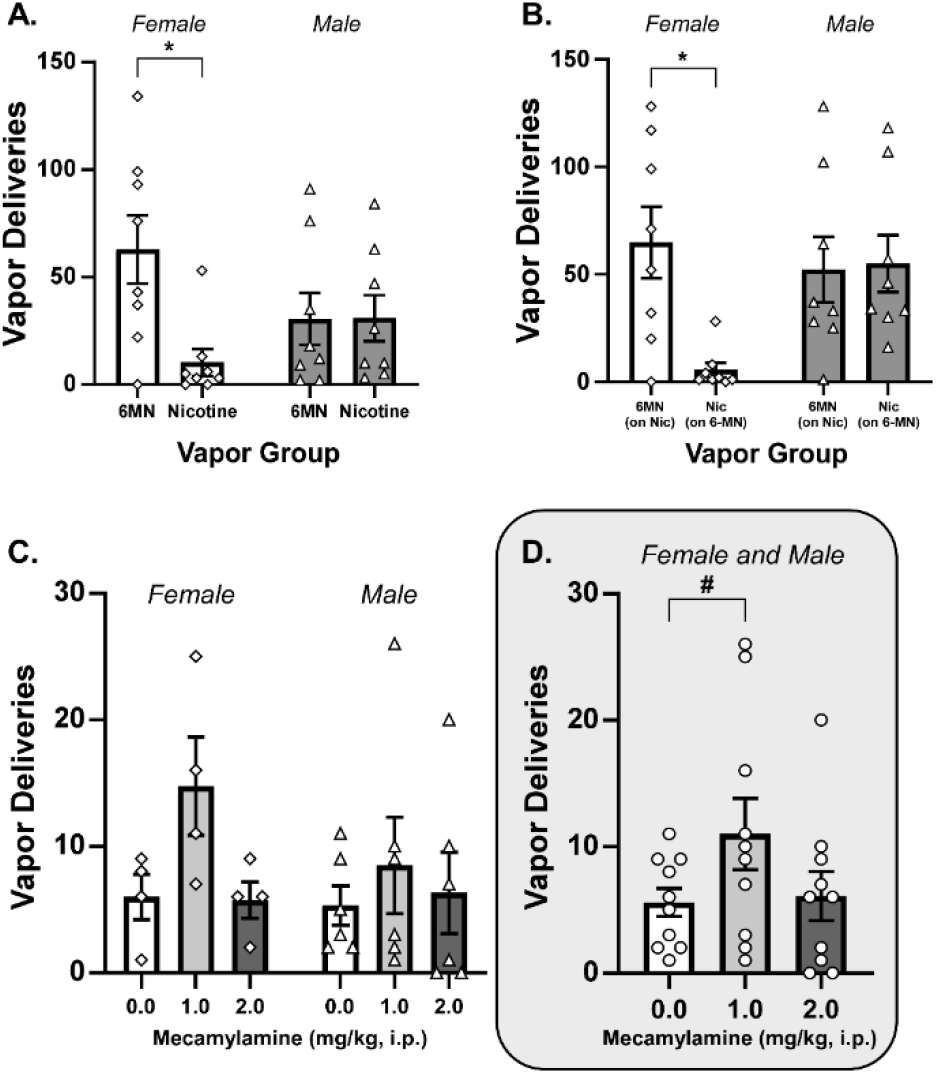
Mean (±SEM) and individual vapor deliveries obtained in 30-minute vapor self-administration sessions. A, B) Sum of deliveries in 9 sessions for each of nicotine (3 mg/mL) and 6-MN (3 mg/mL) in a crossover design. C, D) 6-MN (3 mg/mL) vapor deliveries obtained in individual sessions following pre-treatment with saline or mecamylamine (1, 2 mg/kg, i.p.). A significant difference between groups is indicated with * and between pre-treatment conditions with #.

Due to the initial results in Group 1, only the 0.0 and 1.0 mecamylamine conditions were examined in the swapped drug condition and there were no effects of pre-treatment condition confirmed. There was, however, a significant effect of mecamylamine pre-treatment condition in the male and female groups originally trained on 6-MN and evaluated on 6-MN (**Figure 5C,D**). For this analysis, rats who did not obtain at least one reinforcer after vehicle injection were excluded (N=2 male 6-MN; N=4 female 6-MN; N=4 female Nic). Preliminary analysis including all four groups confirmed a significant interaction between Group and Pre-treatment condition (F (4.871, 29.23) = 3.44; P=0.05) and analysis limited to the groups trained on 6-MN confirmed a significant effect of Pre-Treatment (F (2, 16) = 4.54; P=0.05), but not of Sex or the interaction of Sex with Pre-Treatment. The post-hoc analysis of the marginal mean (i.e., across sex) confirmed a significant difference between vehicle and 1.0 mg/kg mecamylamine.

### 3.5 Experiment 5: Effect of nicotine and 6-methyl nicotine on wheel activity

#### Cohort 1

Wheel activity was suppressed by injection with nicotine or 6-MN (**Figure 6**), with some differences observed across compounds at ~equivalent doses of 0.2 nicotine tartrate / 0.075 mg/kg 6-MN freebase.

**Figure 6.**
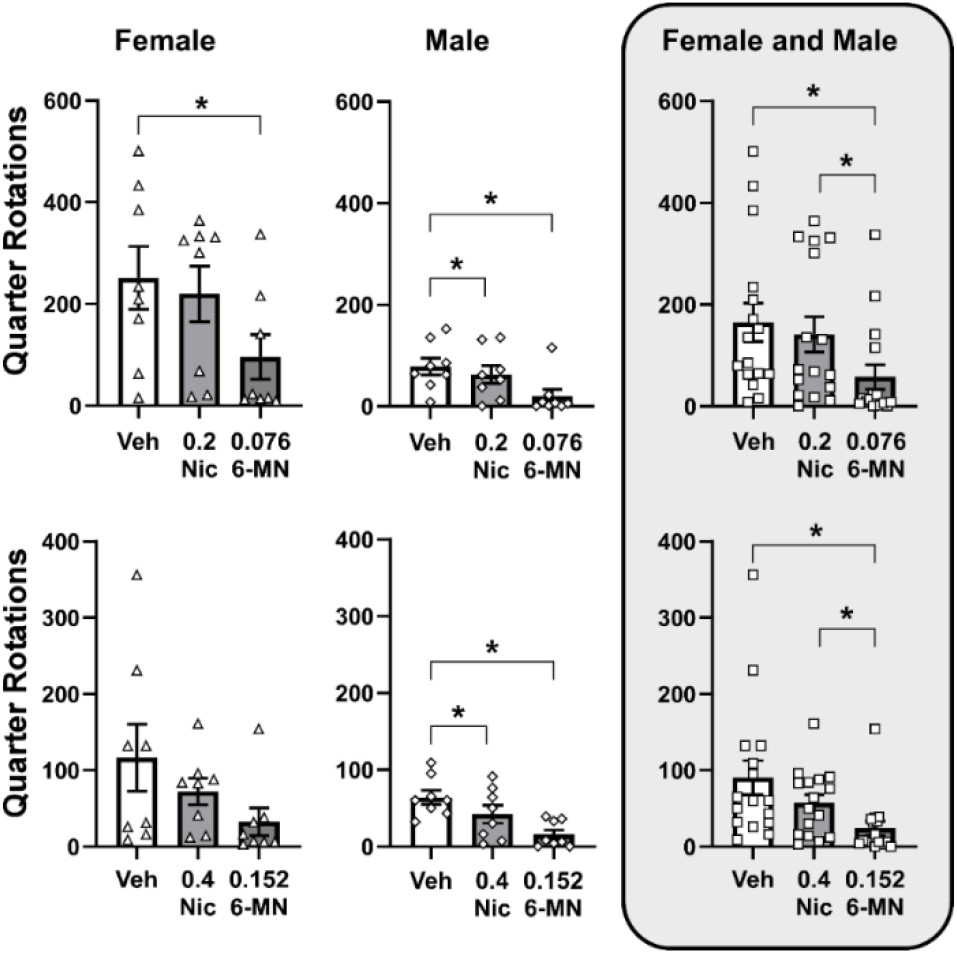
Mean (±SEM; N=8 per sex) and individual session wheel activity following injection with vehicle, (-)-nicotine ditartrate (0.2, 0.4 mg/kg, i.p) or (-)-6MN freebase (0.076, 0.152 mg/kg, i.p.) for Cohort 1. A significant difference from the vehicle condition is indicated with *.

The two-factor analysis of the lower dose experiment confirmed significant effects of Sex (F (1, 14) = 6.91; P<0.05) and of Drug Condition (F (1.605, 22.48) = 16.71; P<0.0001) on wheel activity and analysis of the higher dose experiment confirmed a significant effect of Drug Condition (F (1.434, 20.07) = 7.50; P<0.01) on activity. The Tukey post-hoc tests confirmed significant reductions after the 6-MN injection relative to vehicle or nicotine injection for both higher and lower doses, collapsed across sex. Within the female group the 0.076 mg/kg 6-MN injection reduced activity relative to the vehicle, whereas within the male group, activity was lower after either nicotine or 6-MN in each dose condition compared with the respective vehicle injection.

#### Cohort 2

Due to unforeseen complications we completed only N=4 per sex in the vehicle condition and N=6 per sex in the nicotine (0.2 mg/kg, i.p.) and 6-MN (0.076 mg/kg, i.p.) conditions out of N=8 per sex total. The mixed-effects analysis of summed wheel activity confirmed significant effects of Sex [F (1, 14) = 6.60; P<0.05] and Drug Condition [F (0.3444, 2.066) = 40.35; P<0.05] and the Tukey post-hoc test of the marginal mean confirmed fewer rotations after 6-MN compared with either vehicle or nicotine injection (**Figure 7**). The Sidak post-hoc test further confirmed that males engaged in significantly less wheel activity after vehicle injection.

**Figure 7.**
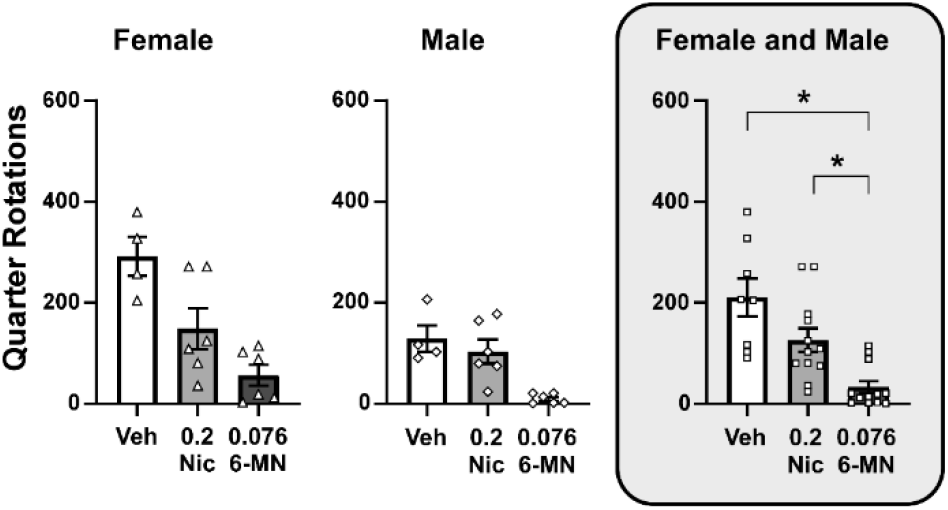
Mean (±SEM) and individual session wheel activity following injection with vehicle, (-)-nicotine ditartrate (0.2 mg/kg, i.p) or (-)-6MN freebase (0.076 mg/kg, i.p.) for Cohort 2. A significant difference from the vehicle condition is indicated with *.

## 4. Discussion

This study found that the inhalation of (-)-6-methyl nicotine (6-MN) produced anti-nociceptive and thermoregulatory effects similar to those produced by the inhalation of (-)-nicotine in rats. Body temperature was lowered, as was previously shown for nicotine inhalation (Javadi-Paydar et al., 2019), and warm-water tail withdrawal latency was modestly increased. The thermoregulatory effect of 6-MN was dose-dependent in the Group 1 female rats across the 1-30 mg/mL concentrations, with a less dose-dependent impact in male rats. In the direct comparison study in Group 2, the impact of inhaled 6-MN and nicotine on body temperature and nociception was similar across 30 and 60 mg/mL concentrations, save that the higher concentrations produced a longer lasting hypothermia in male rats. Self-administration of vapor was equivalent for nicotine and 6-MN in the male rats, and was similar to the males for females initially trained on 6-MN. Pre-treatment with the nicotinic receptor antagonist mecamylamine significantly increased responding for 6-MN vapor.

The observed maximum effect on temperature of ~1°C is consistent with prior observations for nicotine with body temperature assessed telemetrically (Javadi-Paydar et al., 2019; Javadi-Paydar et al., 2024), and with observations that nicotine 0.08-1.0 mg/kg injected parenterally reduces rat body temperature by 1-1.5C (Dilsaver and Davidson, 1987; Dilsaver et al., 1988; Javadi-Paydar et al., 2019). The antagonist mecamylamine attenuated the reduction of body temperature after nicotine inhalation even while producing a slight reduction of body temperature after PG inhalation. This is partially consistent with a prior result where mecamylamine produced small reductions in intraperitoneal temperature (Javadi-Paydar et al., 2024), albeit in that case mecamylamine also potentiated the impact of nicotine to reduce body temperature. 6-MN did not alter temperature significantly after either saline or mecamylamine pre-treatment in this study, which may reflect some degree of tolerance due to the sequence of experiments in the group.

The anti-nociceptive effects of nicotine and 6-MN inhalation in this study were consistent with those reported in middle aged female rats for 6-MN injection (Taffe et al., 2026), on the order of what has been observed after nicotine or THC vapor inhalation in rats (Nguyen et al., 2016; Ogden et al., 2026), but of relatively small magnitude compared with, e.g., opioid vapor inhalation (Gutierrez et al., 2022; Nguyen et al., 2019). The antinociceptive effects of *injected* nicotine are also of a magnitude less than can be produced with an opioid in mice (Berrendero et al., 2002; Semenova et al., 2012) and rats (Carstens et al., 2001; Hunt et al., 1998; Khan et al., 1998).

The vapor self-administration experiment found that reinforcement was variable across individuals, possibly due to selecting the relatively low 3 mg/mL training concentration. No reliable sex differences for 6-MN self-administration were found, although the female animals trained initially on nicotine had a high percentage of long term refusers. Relatedly, female Sprague Dawley rats self-administered fewer *intravenous* infusions of nicotine than did Sprague Dawley males, whereas Long-Evans rats did not differ by sex (Leyrer-Jackson et al., 2021). We previously showed female Sprague Dawley rats exposed as adolescents to repeated non-contingent nicotine vapor self-administer more vapor deliveries than males at a 10 mg/mL concentration (Gutierrez et al., 2024b), suggesting some complexity in the relationship of sex and strain to the self-administration of nicotinic drugs. Although the male groups in the present study self-administered more vapor deliveries in the second 9 sessions compared with the first 9 sessions, this did not vary by the initial drug identity and the males self-administered comparable amounts of nicotine and 6-MN vapor within the first and second set of sessions. Importantly, the administration of a moderate dose of the nicotinic antagonist mecamylamine increased the number of 6-MN vapor deliveries in both sexes, supporting the inference of pharmacological specificity of reinforcement.

Wheel activity was significantly reduced by injection of (-)-nicotine ditartrate or (±)-6MN ditartrate injection in middle aged female Wistar rats in our prior report (Taffe et al., 2026), with similar effects at the 0.8 mg/kg dose of each compound. Wheel activity was similarly suppressed by nicotine vapor inhalation in a prior study (Gutierrez et al., 2024a). Here, we confirm the effects of (-)-6MN freebase injection in male and female Sprague Dawley rats, with only a minor effect of 0.2-0.4 mg/kg (-)-nicotine ditartrate injection limited to male rats. This difference, observed after molecular weight equivalent doses of the (-) isomers, suggest potentially enhanced potency of the 6-MN over nicotine.

Sex differences in the impact of 6-MN and nicotine in this study were minimal despite significant differences between male and female rats in terms of basal body temperature and voluntary wheel activity. One exception is that the temperature of the male rats stayed low through the 60-minute observation after drug inhalation, whereas the female rats’ temperature returned to baseline at this time point. This difference may be due to the larger body size of the male rats who were 412.4 g (Stdev 23.1) compared with 245.9 g (Stdev 10.5) at the start of the dose-effect evaluation. Female Sprague Dawley rats (4-5 months) are more sensitive to the antinociceptive effects of nicotine after i.c.v. injection (Craft and Milholland, 1998), but no sex difference was observed here, possibly due to the limited effect size.

This study used the more active (-) isomers of nicotine and of 6-MN. However, given that 6-MN may be synthetized in some commercial preparations, there may be need for future work comparing the racemate with the more active isomer. While some evidence suggests the (+)-nicotine isomer is more than 100 fold less potent (Abood et al., 1978), other studies indicate it is only 9 fold less potent (Meltzer et al., 1980). The two isomers were equipotent in evoking responses from frog olfactory neurons (Thurauf et al., 1995), and our prior work suggested ~equipotency of the 6-MN racemate with (-)-nicotine in rats (Taffe et al., 2026).

Our initial report described the effects of 6-MN in highly nicotine-experienced middle aged female Wistar rats (Taffe et al., 2026). Here, we show that those effects generalize to both sexes of young adult animals, of a different strain, that have not had extensive prior nicotine exposure. A comparison of the thermoregulatory impact of 6-MN (5, 10, 30 mg/mL) in young adult females in the present work and in the middle aged females in the prior study suggest a modest tolerance in the latter. Alternately, this may be due to a different body size or the fact that Wister rats may be less sensitive than Sprague Dawley rats to hypothermia induced by some drugs, e.g., as demonstrated for THC (Taffe et al., 2021).

Overall, the roughly similar effects of nicotine and 6-MN at the same e-liquid concentrations in this study contrast somewhat with an inference of significantly higher 6-MN potency based on analysis of ~3-6 mg of 6-MN per gram of e-liquid in several commercial products that were labeled as containing 50 mg/g, presumably as a cue to the nicotine-adapted consumer (Erythropel et al., 2024; O’Connor et al., 2026). The inference of roughly equivalent potency is, however, similar to a prior report of similar potency of nicotine and 6-MN in a drug-discrimination assay in rats (Meltzer et al., 1980).

## Acknowledgements

The authors thank Christianne J Perral, for technical assistance. The authors thank Wayne Mascarella, Ph.D. for assistance with sourcing and verifying the 6-methyl nicotine used in these investigations. These studies were supported by the Tobacco Related Disease Research Program (T33IR6653 MAT) and the US National Institutes of Health (R01 DA057423). The funding entities had no influence on the study design, data interpretation, manuscript creation or in the decision of when and what to publish from the studies conducted.

## Contributors MAT

Conceptualization; Data curation; Formal analysis; Funding acquisition; Methodology; Project administration; Resources; Supervision; Visualization; Roles/Writing - original draft; Writing - review & editing. HSK: Investigation; Writing - review & editing. TAD: Investigation; Writing - review & editing. TRC: Investigation; Writing - review & editing. SRMU: Investigation; Writing - review & editing. YG: Investigation; Methodology; Writing - review & editing. SAV: Investigation; Methodology; Writing - review & editing. All authors reviewed the manuscript.

